# Unravelling Key Interactions and the Mechanism of Demethylation during hAGT mediated DNA Repair via Simulations

**DOI:** 10.1101/2022.07.26.501539

**Authors:** Shruti T G, Shakir Ali Siddiqui, Kshatresh Dutta Dubey

**Author notes:** Authors have Equal Contributions (Shruti T G, Shakir Ali Siddiqui).

## Abstract

Alkylating agents possess the biggest threat to the genomic integrity of cell by damaging DNA bases through regular alkylation. Such damages are repaired by several automated machinery inside cell. O6-alkylguanine-DNA alkyltransferase (AGT) is such an enzyme which performs the direct repair of an alkylated guanine base by transferring the alkyl group to a Cysteine residue. In the present study using extensive MD simulations and hybrid QM/MM calculations, we have investigated the key interactions between the DNA lesion and the hAGT enzyme and elucidated the mechanisms of the demethylation of the guanine base. Our simulation shows that the DNA lesion is electrostatically stabilized by the enzyme and the Arg135 of hAGT enzyme provides the main driving force to flip the damaged base into the enzyme. The QM/MM calculations show demethylation of damaged base as a three step in thermodynamically feasible and irreversible manner. Our calculations show that the final products forms via Tyr114 in a facile way in contrast to the previously proposed Lys-mediated route.

## INTRODUCTION

The genomic integrity of a cell is constantly under threat by some extracellular and intracellular chemicals which can damage the nucleotide base of a DNA by covalently attaching an alkyl group^1–6^. Such damaged DNA can cause deleterious mutations and cytotoxicity in cells^2,3,6–10^, and therefore, the cell has the ultimate machinery to repair such DNA damage. This repairing machinery is mostly done by some proteins and/or enzymes that have evolved particularly for this purpose, mainly via three different mechanisms: a) Photolesions through photolyases by UV induction b) reversal by O6-alkylguanine-DNA alkyltransferase (AGTs) and c) reversal by AlkB family dioxygenases^1,3^. However, the last two mechanisms, i.e., the damage reversal by AGT and AlkB enzymes are a direct DNA repair mechanism through the de-alkylation of damaged base and are, hence, believed to be the most efficient way to repair DNA lesions^1^. It is anticipated that the mechanism of the repairing is highly correlated with the position of the alkylation attack^11^. For example, if alkylation occurs on the N^7^ position of the guanine base, it results in an innocuous lesion that is mostly repaired through depurination. Similarly, when alkylation occurs at the N^3^ position of the nucleotide base, the resulting lesion blocks DNA replication and is repaired by AlkA or AlkB proteins. The third vulnerable site for alkylation is the oxygen atom (O^6^) of the DNA base to produce the O^6^-methylaguanine^6,7^(see Figure 1a). This lesion is believed to be highly mutagenic and it mispairs with thymine to produce a transition mutation of G: C →T: A during DNA replication^1,10,12,13^. These lesions are repaired by the O6-alkylguanine-DNA alkyltransferase (AGT) family of proteins by a suicidal direct repairing mechanism^1,2,6,8–10,14–16^. In the current work, we have highlighted the key interactions and the mechanism of the DNA repairing of human AGT protein using MD simulations and hybrid QM/MM calculations.

**Figure 1.**
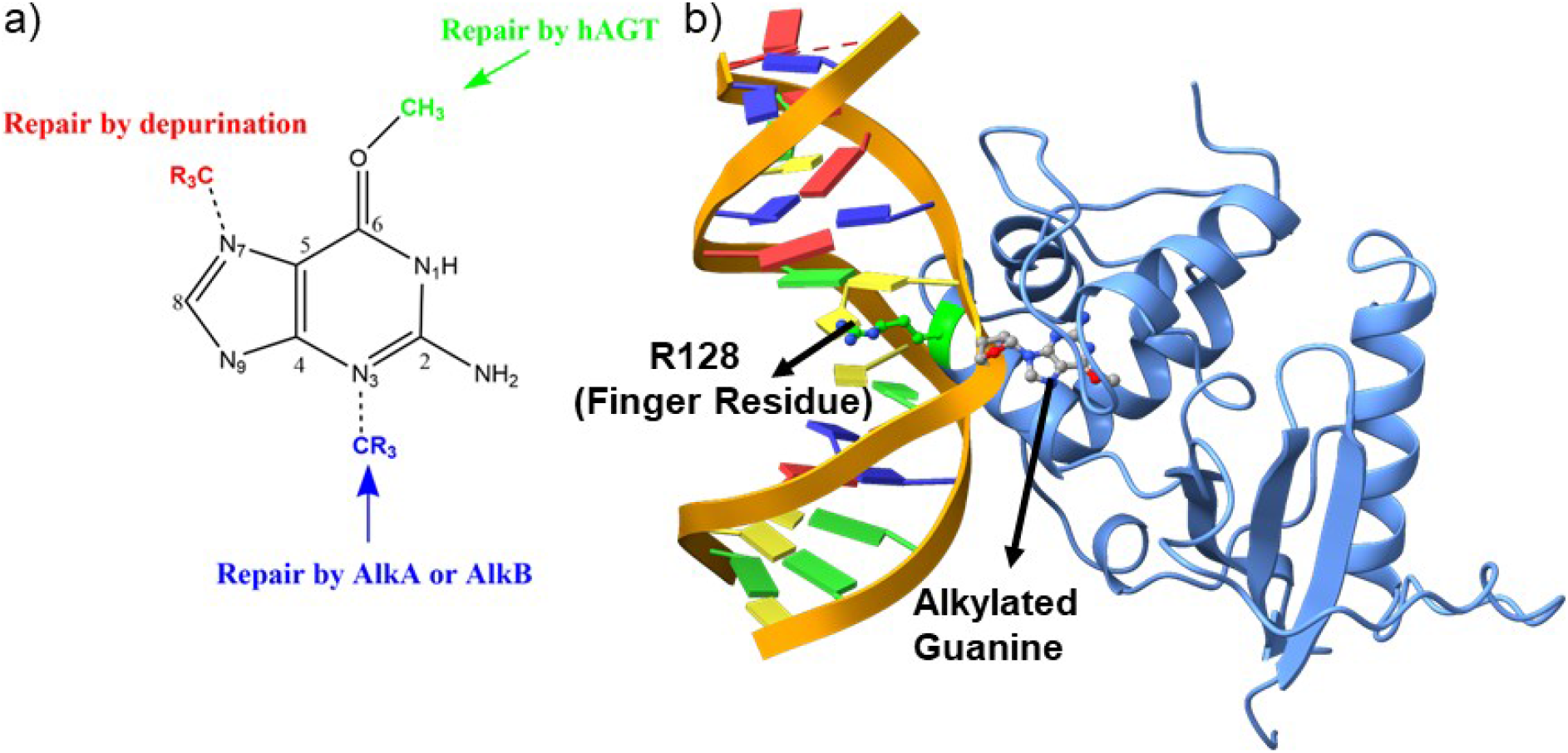
a) DNA repair mechanism based on position of the alkylation b) Structure of the flipped alkylated guanine in hAGT (pdb id 1T38)

The crystal structure of the hAGT protein complex with double-strand DNA (dsDNA) carrying the damaged guanine^3^ provides crucial insight into the base flipping after alkylation. This structure shows the base flipping by a minor grove of the dsDNA^3^ and the modified base is inserted into the active site of the protein^3,9^, which is surrounded by Cys145, Tyr114, Pro140, Ser159, and Tyr158^2,10,15^. Here Cys 145 participates directly in the repair by accepting the alkyl group^2,3,9,10^ while Tyr114 is believed to facilitate a proton transfer in the reaction. An Arg128 residue assigned as the ‘finger residue’ was found to be inserted inside the DNA duplex, in place of the damaged, extrahelical base^2,3,6,9^. Interestingly, the role of this ‘finger residue’ is supposed to be instrumental in identifying the damaged base^13^ by sliding over DNA bases^15^ by checking the weakened base-base interaction due to the alkylation of the guanine base (sometimes thymine).

The experimental structure of hAGT with the damaged DNA provided a good starting geometry to validate the repair mechanism using computational tools. However, unlike AlkB where the mechanism of the DNA repair has been extensively studied^1^, the repair mechanism by hAGT is relatively less elucidated. Jena et al^17^ performed DFT only study to explore the repair mechanism by hAGT and proposed a three-step pathway for the repair mechanism. According to their study, in the first step, deprotonation of Cys145 occurs via a water-mediated mechanism from His146. In the second step, protonation at the N3 position takes place via Tyr114 and in the last step demethylation of guanine occurs through Cys145^2,15^. However, the proposed reaction profile was thermodynamically not feasible since they conducted a DFT only study without the inclusion of the protein and DNA molecules. Another study by Hou et al^18^ used a more accurate QM/MM method to explore the repair mechanism of this enzyme. Interestingly, this study shows that the methyl transfer from damaged DNA to cysteine is a reversible process, however, it is well known that the DNA repair by hAGT is suicidal and irreversible. We, therefore, planned to re-investigate the mechanism of the repair by hAGT using extensive MD simulations and hybrid QM/MM calculations. In the present study, we have used comprehensive MD simulation of hAGT enzyme with dsDNA bearing an alkylated guanine to study the interactions between the enzyme and the modified base and, performed hybrid QM/MM calculations to validate the reaction mechanism of the direct DNA repair by hAGT.

## 2. COMPUTATIONAL DETAILS

We have performed the MD simulations to study the conformational changes and protein-DNA interactions while hybrid QM/MM calculations were performed for the reaction mechanism. The details of each calculation are discussed below.

### 2.1 System Setup

The initial coordinate of the hAGT in complex with dsDNA was imported from the protein data bank (PDB id: 1T38)^3^. The crystal structure contains an alkylated guanine base flanged out from the DNA strand and was buried inside the protein site. The parameters for the modified base were prepared using an antechamber module of the Amber MD program of a QM optimized geometry at HF/6-31 g(d,p) level of theory. For protein, we used Amber ff19SB^19^ forcefield while for DNA we used refined Barcelona forcefield implemented in the Amber MD library. A few Na+ ions were added to the protein surfaces to neutralize the total charge of the system depending upon the charge of each complex prepared separately. Finally, the resulting systems were solvated in an octahedral box of OPC water model each extended up to a minimum cut-off of 10 Å from the protein boundary.

### 2.2 MD Simulations

After proper parametrization of the system, to remove bad contacts, minimization was performed in two stages using a combination of steepest descent (5000 steps) and conjugate gradient (5000 steps) methods. In the first stage water position and conformations are relaxed keeping the protein fixed. Thereafter the whole complex was minimized. Subsequently, the system is gently annealed up to 300K under the NVT ensemble for 50ps. After that, 1ns of density equilibration was performed under an NPT ensemble at a target temperature of 300K and pressure of 1 atm by using Langevin thermostat^20^ and Berendsen barostat^21^ with a collision frequency of 2ps and pressure relaxation time of 1ps. This 1 ns density equilibration is a weakly restrained MD simulation in which the system is slowly released to achieve uniform density after heating under periodic boundary conditions. Then after we remove all the restraints applied before and the system gets equilibrated for 3 ns to get a well-settled pressure and temperature for chemical and conformational analyses. Thereafter, a productive MD simulation was performed using the Monte Carlo barostat^22^ for 100 ns for each complex system.. Therefore, we performed a total of 300 ns simulations including all three replicas for each system. During all the MD simulations, covalent bonds containing hydrogens were constrained using the SHAKE^23^ algorithm, and Particle Mesh Ewald (PME)^24^ method was used to treat long-range electrostatic interactions with the cut-off set as 10 Å. All MD simulation was performed with the GPU version of the AMBER 20 package^25^. The MD trajectory analysis was done with the CPPTRAJ^26^ module of AMBER20. The visualization of the MD trajectories was performed by VMD^27^. The binding free energy was calculated using Molecular Mechanics Generalized Born Surface Area method (MMGBSA), the details and other applications for the nucleic acid complexes have been discussed elsewhere.^28–29^

### 2.3. QM/MM Methodology

The mechanism of reaction during the base-repairing was calculated using hybrid QM/MM calculations for the representative snapshots from MD simulations. The active region in QM/MM calculations in all the systems involves the protein residues and water molecules present within the cutoff of 8Å from the active oxidant heme. The atoms in the selected “active region” (mainly from the MM part) interact with the QM zone through electrostatic and van der Waals interactions and the corresponding polarization effects were considered in the subsequent QM/MM calculations. All QM/MM calculations were performed with ChemShell^30,31^, by combining the Turbomole^32,33^ for the QM part, and DL_POLY^34^ for the MM part. The MM part was described using ff19SB forcefield. To account for the polarizing effect of the protein environment on the QM region, an electronic embedding scheme was used. Hydrogen link atoms with the charge shift model were employed for treating QM/MM boundary. During QM/MM geometry optimizations, potential energy surface scanning, and frequency calculations, the QM region was treated using the hybrid B3LYP functional with a def2-SVP basis set. The energies were further corrected with the Grimme dispersion correction. All of the QM/MM transition states were located by relaxed potential energy surface (PES) scans followed by full TS optimizations using the P-RFO^35^ optimizer implemented in the HDLC code.

## 3. RESULTS AND DISCUSSION

### 3.1. MD simulations of hAGT enzyme with the alkylated base after flipping

To study the mechanisms, we started with the MD simulation of the flipped methylated guanine (m-GUA) with hAGT to investigate the conformational changes if any. During the entire course of the simulation of 500 ns time, we didn’t observe much conformational changes in the dsDNA which can be validated by the low RMS (root mean square) deviation as shown in Figure 2 relative to the enzyme. We found that most of the deviation in the enzymatic site comes due to several loop regions of the hAGT enzyme. To further validate the flexibility, we also supplemented our results with the RMSF (root mean square fluctuations) for residues during the simulations.

**Figure 2.**
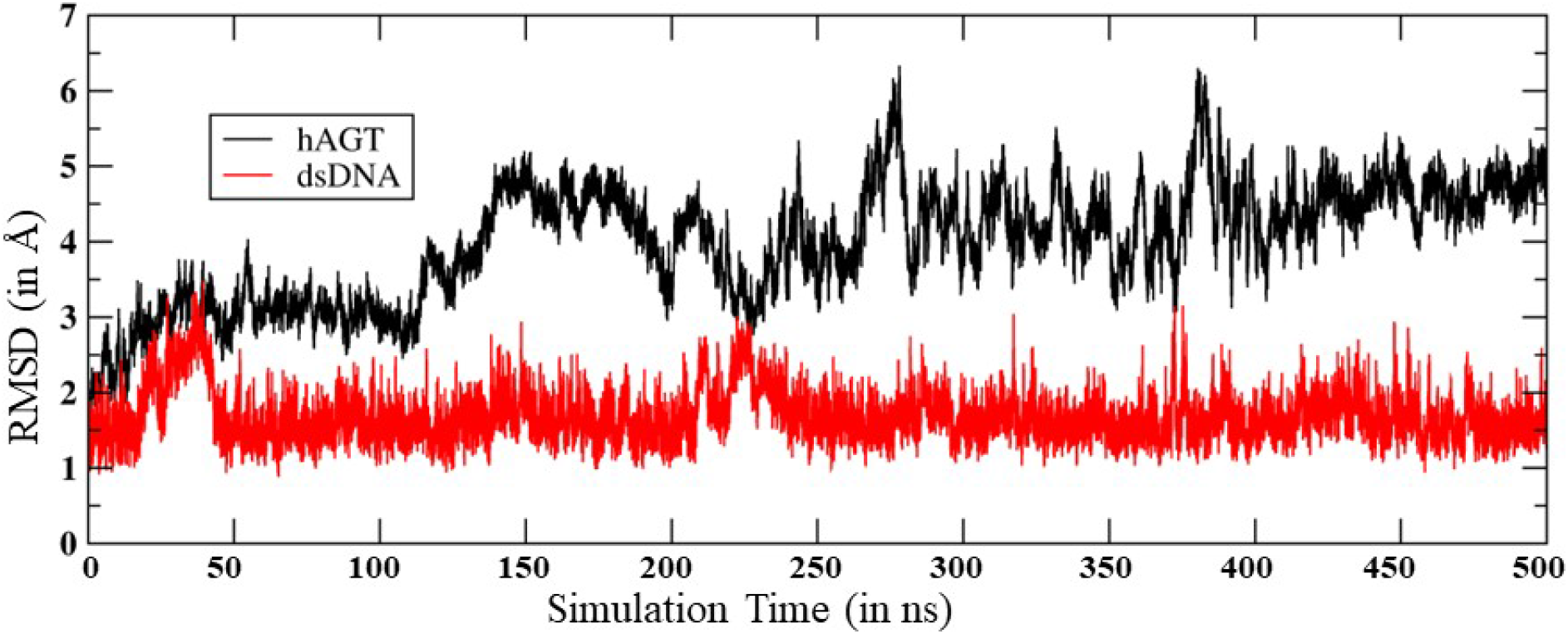
RMS deviation for hAGT and dsDNA during the MD simulations.

As can be seen (Figure 3), the region of the highest flexibility comes from the residues 30-50 which is from the loop region of the hAGT. We note here that, this loop region is the zinc-binding region which may have functional significance in the direct repair of the DNA lesion.^11^ Interestingly, the m-GUA (residue 178) shows very small flexibility relative to other DNA regions, which might be due to the strong binding with the catalytic residues of the hAGT enzyme.

**Figure 3.**
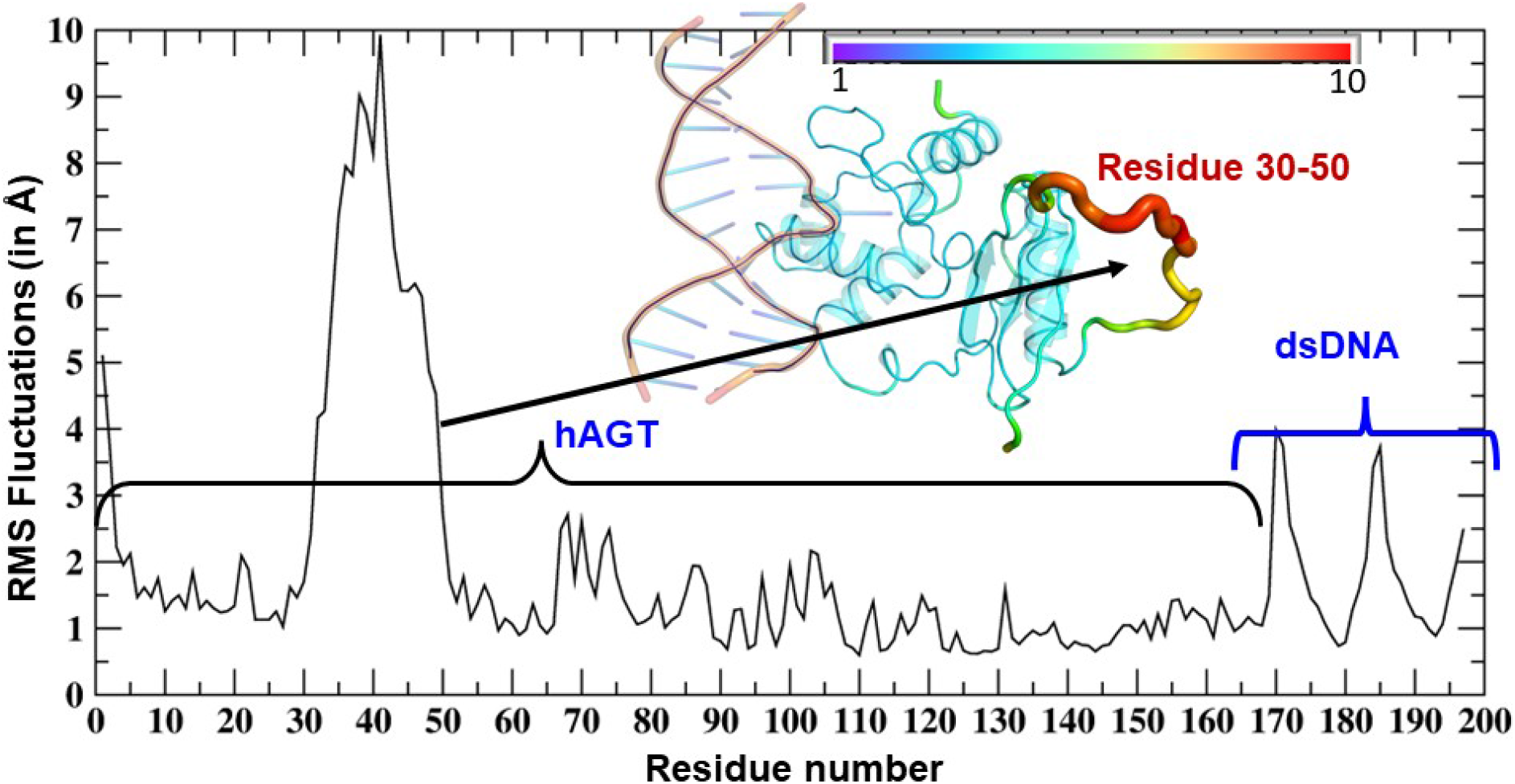
RMSF of dsDNA and hAGT complex. The thickness of the tube in the inset represents the region of the highest flexibility.

A representative snapshot from the MD simulation is shown in Figure 4. As can be seen, R128 (finger residue) occupies the vacant space of the guanine and interacts strongly with the orphaned cytosine (Figure 4a). On the other hand, the flipped m-GUA is well installed in the catalytic site and maintains a rigid conformation throughout the entire simulation (Figure 4b). The C145 which is supposed to abstract the methyl group of the m-GUA resided proximal to the m-GUA and maintains proximity with the methylated end. Interestingly, we found a well-organized water chain connecting His146 to Cys145—Tyr159—Lys165 via WAT1 and WAT2. The role of His146 has already been proposed during the proton transfer from Cys145 during de-methylation of the m-GUA. To quantify the interaction of the m-GUA in the enzymatic site, we calculated the binding free energy of m-GUA into the hAGT using MMGBSA (Molecular Mechanics Generalized Born Surface Area) method (Table 1).

**Figure 4.**
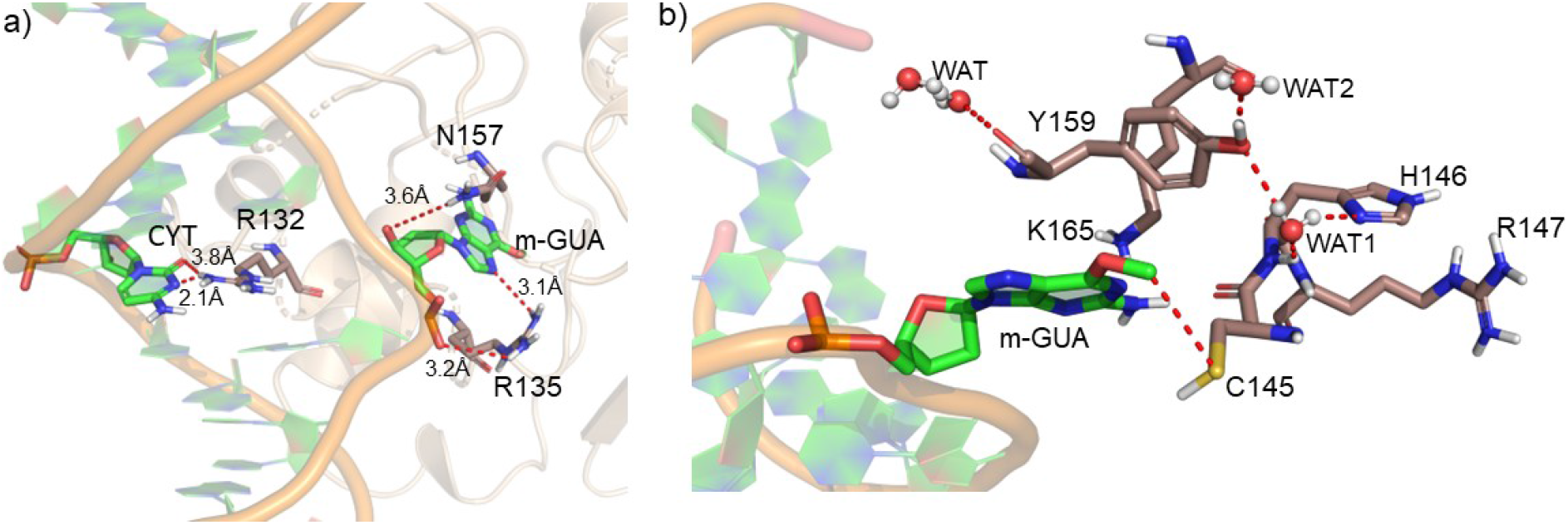
Representative snapshot from the MD simulation a) Interaction of the DNA bases with the protein b) Interaction of the m-GUA with catalytically important residues.

Our calculations show a favorable binding free energy of −32.94 kcal/mol which indicates the twisting of the m-GUA base could be spontaneous and the interactions of the enzymatic site might be the driving force for the twisting.

**Table 1.**
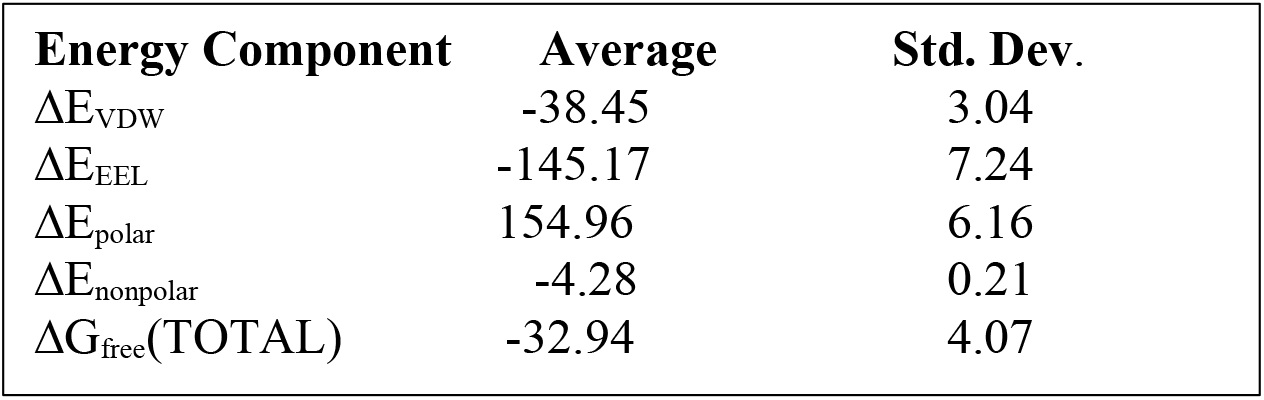
Total binding free energy calculations by MMGBSA method. All values are in kcal/mol.

| Energy Component | Average | Std. Dev. |
| --- | --- | --- |
| $\Delta E_{VDW}$ | -38.45 | 3.04 |
| $\Delta E_{EEL}$ | -145.17 | 7.24 |
| $\Delta E_{polar}$ | 154.96 | 6.16 |
| $\Delta E_{nonpolar}$ | -4.28 | 0.21 |
| $\Delta G_{free}(TOTAL)$ | -32.94 | 4.07 |

Furthermore, to quantify the residue-wise interactions with the m-GUA, we calculated the residue-wise decomposition of the total binding free energy using the MMGBSA method (Figure 5). As can be seen, R135 which is close to the DNA helix, applies the strongest interaction on the m-GUA, and therefore we believe, it could be the driving interaction that might lead to the flipping of the methylated DNA base. Furthermore, the catalytic residues e.g., Tyr114, Cys145, Tyr158, and Lys165 also show significant interactions with the m-GUA. Here, it is quite noteworthy We see that most of the residues which interact with the m-GUA are either polar or charged except Met134 which shows that the binding of the m-GUA in hAGT is predominantly electrostatically driven.

**Figure 5.**
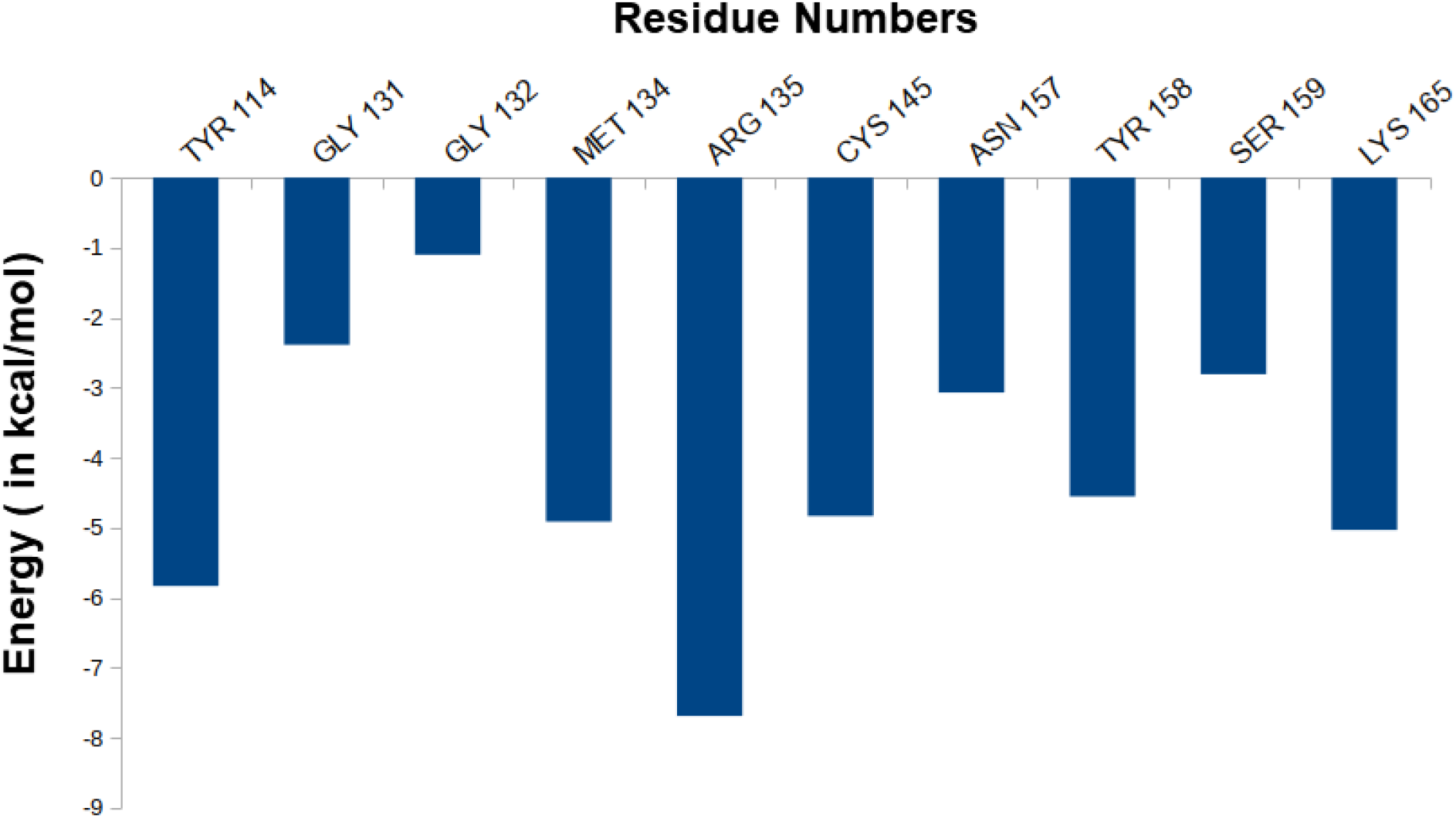
The residue-wise decomposition of the total binding free energy of the m-GUA.

### 3.2. Mechanistic Elucidation of O6-demethylation via QM/MM Calculations

In the previous section, we have seen that the m-GUA is well installed in the catalytic site of the hAGT enzyme, and it is surrounded by catalytically important residues such as Cy145, Tyr114, Ser159, and Lys165. In addition, we also found a well-organized water channel bridging His146 with Cys145 which can assist the deprotonation of the Cysteine. The deprotonation mechanism of the Cysteine via Histidine is well established and therefore, we have focused only on the demethylation of the m-GUA and formation of the guanine since there were discrepancies in the previous investigation of the mechanism (c.f. introduction section). For so doing, we employed the QM/MM calculations on a snapshot generated from the 500ns of MD simulations to study the mechanistic route of DNA repair (O-demethylation of O6-methylguanine) by hAGT. According to the prior studies, Cys145 is first deprotonated by His146 through a water molecule, followed by O-demethylation of O6-methylguanine(O6G) by CysS^-^, therefore, we used a deprotonated Cysteine during the mechanistic elucidation. A proposed mechanism of the de-methylation of m-GUA is shown in Scheme 1. As can be seen, the anionic charge developed at the ‘O’ atom of guanine following O-demethylation by Cys145 seems to be in resonance with the two ‘N’ atoms, as indicated in the second step of the scheme. Furthermore, the Lys165 and Tyr114 are present near the two ‘N’ atoms that can be protonated it to regenerate the repaired guanine, we, therefore, investigated the two protonation pathways for demethylated guanine from both Lys165 and Tyr114.

**Scheme 1.**
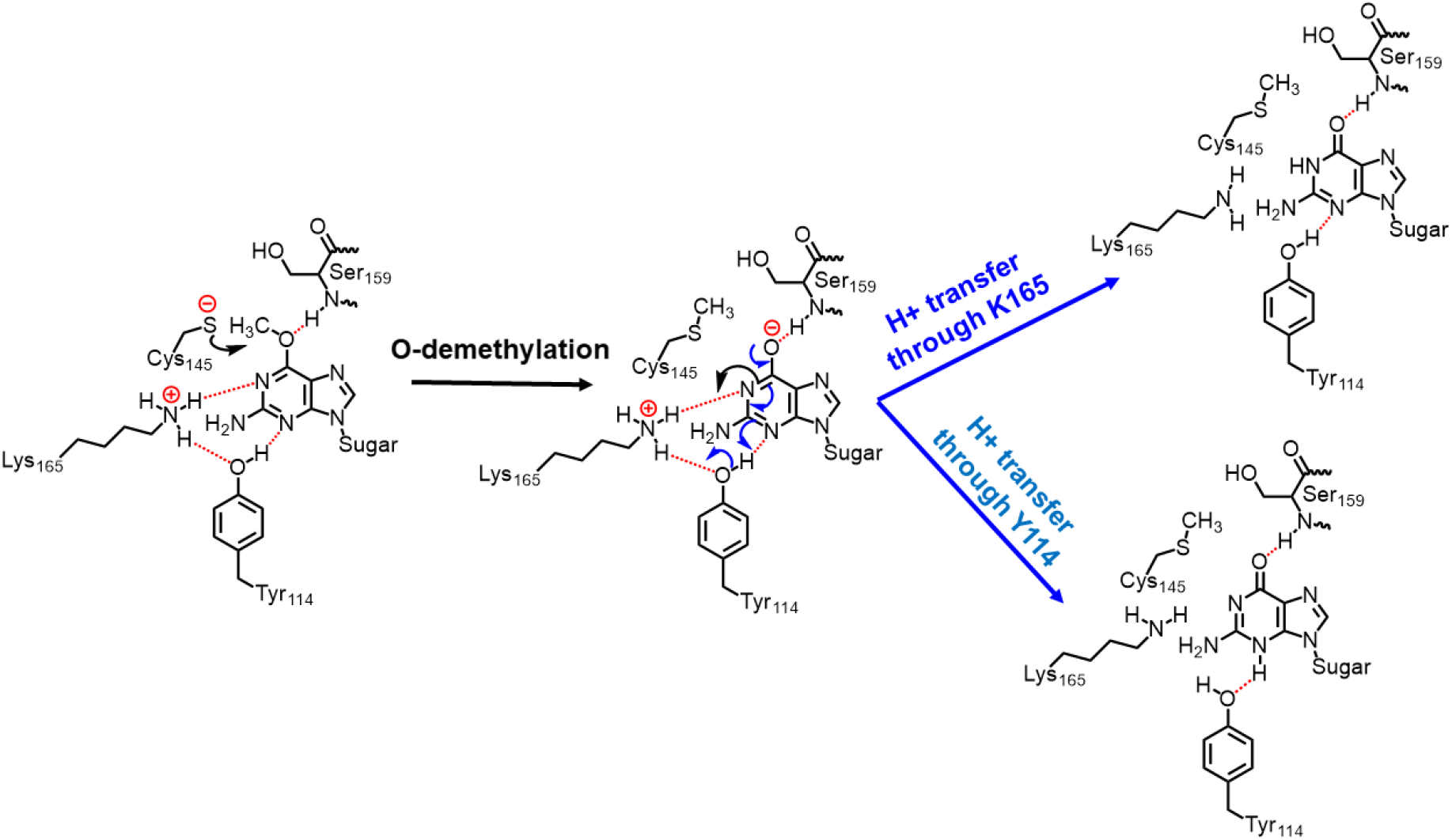
Plausible mechanistic routes for O6-methylguanine repair by AGT. Here, Cys145 anion acts as a nucleophile and Lys165 & Tyr114 act as H^+^ donors.

In order to get a reactant cluster (RC), we picked a representative snapshot from the MD trajectories based on the most populated structure and performed a QM/MM geometry optimization. In the optimized RC, the methyl carbon of the m-GUA was seen to be 3.6 Å distant from the CysS^-^ nucleophile (Figure 6A). To acquire the whole reaction route, we performed the relaxed potential energy surface (PES) scanning, and the reaction profile is shown in Figure 6B. In the first step of the reaction, the CysS^-^ attacks the methyl carbon to perform the O-demethylation of O6-methylguanine through a transition state (TS) barrier of 16.8 kcal/mol (see TS1, Figure 6A), followed by the formation of anionic guanine as intermediate IM.

**Figure 6.**
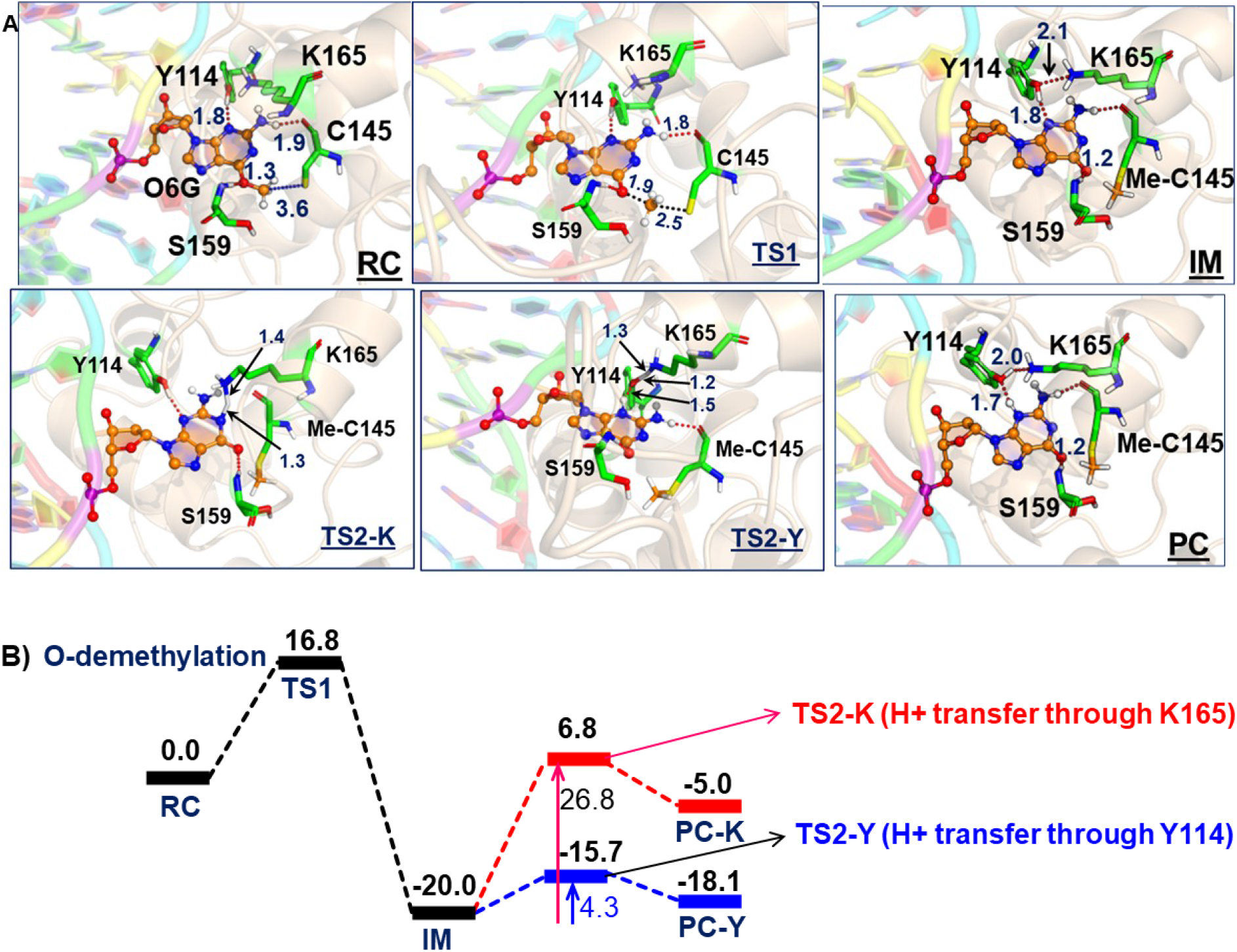
**(A)** The QM/MM optimized geometries along with the key geometric data for RC, TS1, TS2-Y, TS2-K, IM, PC-K and PC-Y. The ‘Me-C145’ represents the methylated Cys145. **(B)** The complete reaction profile diagram. The energy values (reported in kcal/mol) are noted for the optimized structures (B3LYP/def2-SVP) of all the RC, TS, IM, and PC states. All energies are corrected by zero-point energy (ZPE) and Grimme’s dispersion (G-D3).

After the formation of anionic guanine, it needs to be protonated to generate the repaired guanine. As discussed earlier it could be via two pathways: either via the Lys165 or Tyr114 routes. Therefore, we explored both routes through the PES scanning, and the reaction profile for the same is shown in blue and red colors in Figure 6B. The reaction profile in red depicts the relative transition state barrier for H^+^ transfer to guanine via Lys165 while the profile in blue shows the H^+^ transfer to guanine via Tyr114. As can be seen, the TS barrier for H^+^ transfer from Lys165 is observed to be 26.8 kcal/mol which is quite a high barrier for H^+^ transfer reactions in enzymatic reactions. On the other hand, the production of the repaired guanine via H^+^ transfer by Tyr114 is very facile and occurs through a low barrier of 4.3 kcal/mol. Therefore, the second route is relatively preferable over the Lys165 for the protonation of demethylated guanine. Interestingly, we found that as soon as the H^+^ is transferred from Tyr114 onto guanine, one H^+^ is retrieved from Lys165 to rejuvenate itself. Furthermore, we found that the Ser159 which is close to the m-GUA stabilizes the TS and plays a crucial role in the catalysis.

In a nutshell, we can state that this entire DNA repair mechanism (demethylation of O6-methylguanine) consists of three steps: first, Cys145 is deprotonated to function as a nucleophile, then it performs the O-demethylation, and finally, H^+^ transfer occurs from Tyr114 to form the repaired guanine.

**Scheme 2.**
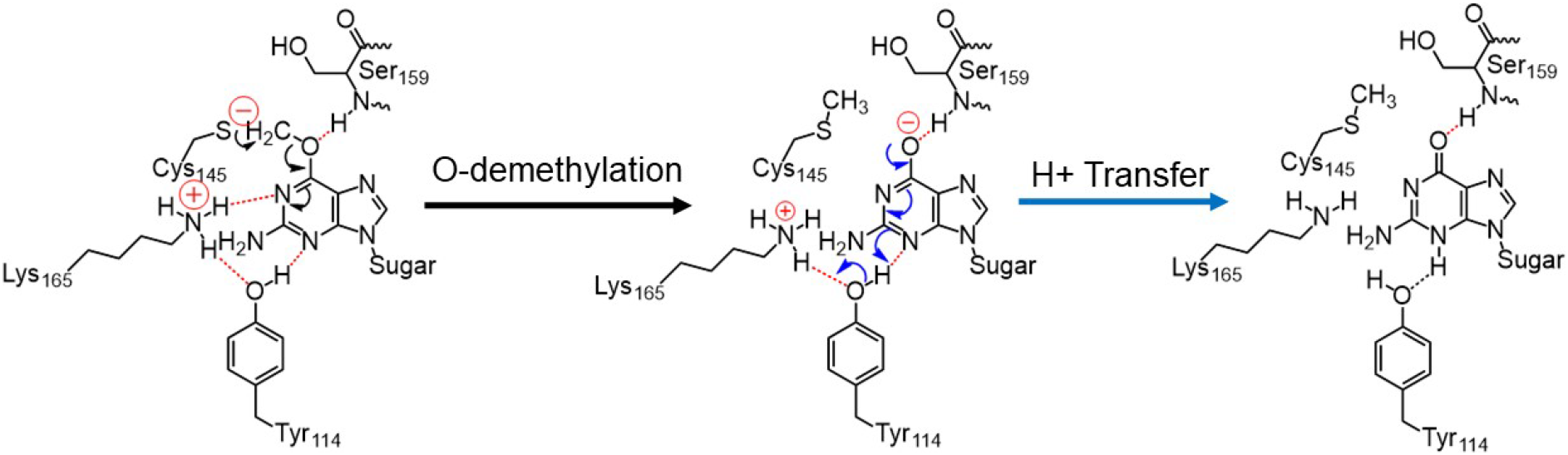
The final mechanism of direct repair of the m-GUA by hAGT.

## CONCLUSIONS

In the present study using comprehensive MD simulation of the double stranded DNA in complex with hAGT enzyme and hybrid QM/MM calculations, we have studied the mechanism of the direct DNA repair by hAGT enzyme. Our MD simulations shows that methylated guanine has several favorable interactions by the protein residues, particularly, Arg135 that provides the driving force for the base flipping. Furthermore, flipped base is thermodynamically stabilized by several polar and charged residues in the active site. The QM/MM study reveals the mechanisms of the demethylation by Cys145 residue and we show that a complete repairing of the guanine can be formed via the Tyr114 rather than Lys165. In addition, our reaction profile shows an irreversible repairing which is in good agreement with the proposed suicidal and irreversible repairing by hAGT enzymes.

## ACKNOWLEDGEMENTS

KDD acknowledges Department of Biotechnology, Ministry of Science and Technology, Govt. of India for Ramalingaswami Re-entry research grant (BT/RLF/Re-entry/10/2017). STG acknowledges Shiv Nadar University for OUR (opportunity for undergraduate research) scheme and she is thankful to Ms. Shalini Yadav, Dr. Surajit Kalita and Ms. Vandana Kardam for their helps and discussions.

## Supporting Information

### H-Bonds Analysis between DNA and protein

H bonds were calculated by uploading 200ns trajectory in VMD, here 10 ps is one frame.

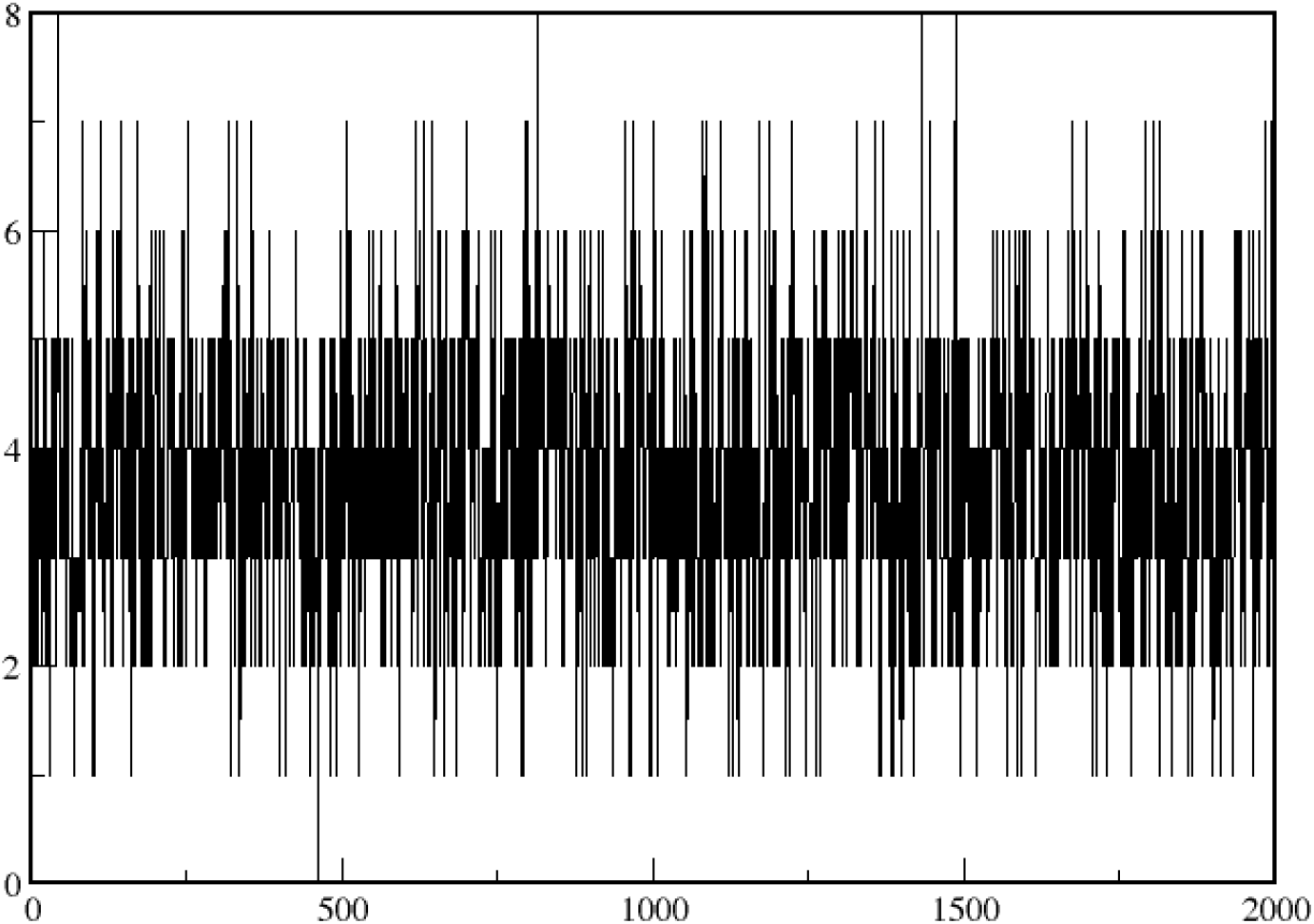

Figure depicts the H-bond analysis for DNA-Protein interaction, with no solvent Consideration. This was performed over 200 ns of simulation time. There is high density of 3/4/5 (maximum 4, thickest band on graph) Hydrogen bonds between DNA and the protein regions of the complex. Some of the major interactions are –

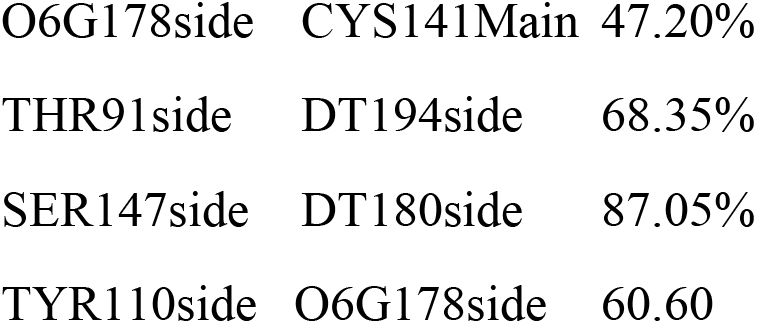

These interactions were also proposed to play significant role in the stability of the extra helical damaged base and prevent its untwisting.

### H Bond between Cys145 and damaged Guanine

H bonds were calculated by uploading 200ns trajectory in VMD, here 10 ps is one frame.

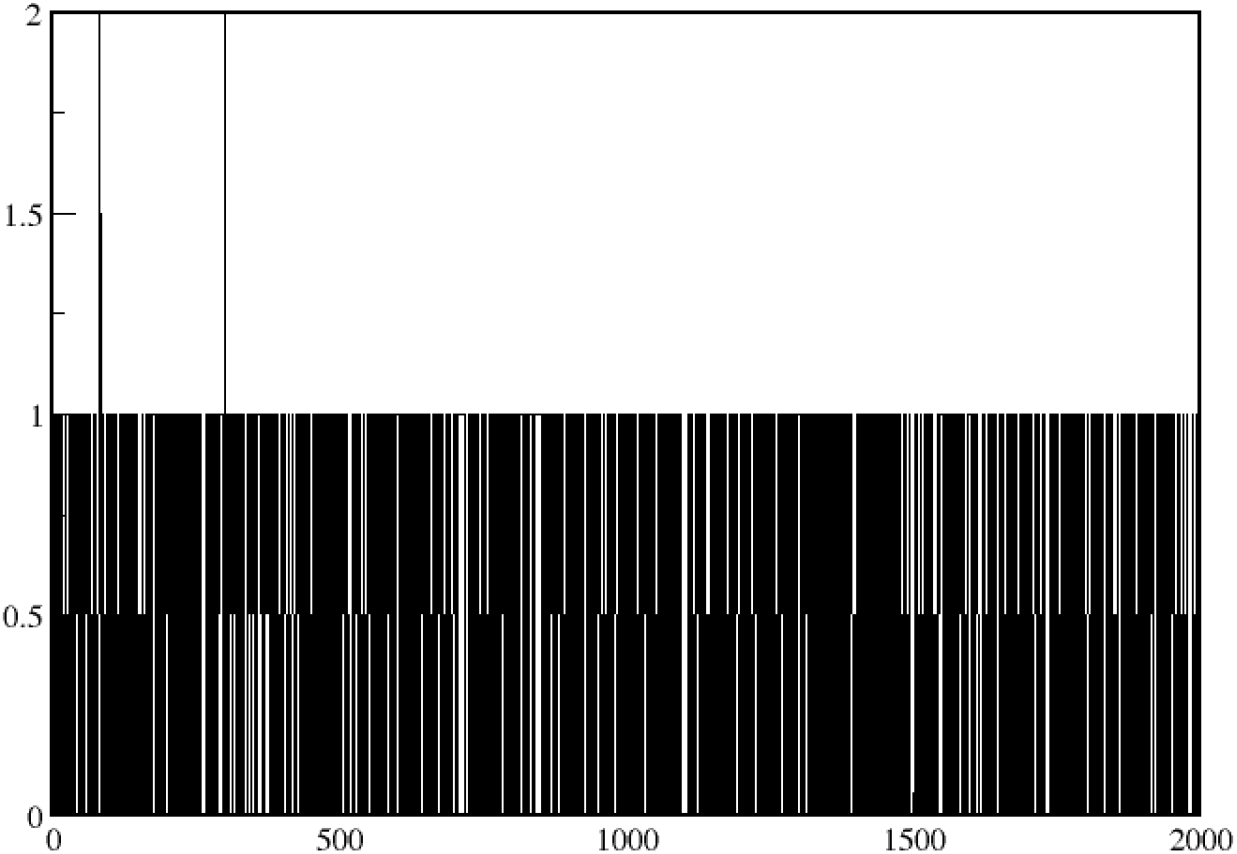

Between the active site residue – Cys145 and the dna, there is almost no instance where HB formations don’t take place. There are HB bonds formed over the entire 200ns, indicating a strong affinity between the O6G substrate and Cys residue-both for ensuring the modified base stays in vicinity of the active site, as well as to mediate the alkyl removal by nucleophilic attack.

### QM Geometry for QM/MM Optimized Structures

#### RC

C −11.6745291 0.0857268 11.8147208

H −12.0679311 −0.0963382 12.8259640

H −12.5189263 −0.0220033 11.1110911

C −10.5880143 −0.9235625 11.4889035

C −9.9845242 −1.6938986 12.4965413

H −10.3439347 −1.5962578 13.5238668

C −8.9354111 −2.5775375 12.2231387

H −8.4796193 −3.1630228 13.0248418

C −8.4626044 −2.7206328 10.9144737

O −7.3930895 −3.5434831 10.6617496

H −7.4011547 −3.8190238 9.7112953

C −9.1043535 −2.0161359 9.8834431

H −8.7863385 −2.1764468 8.8517917

C −10.1494682 −1.1370587 10.1719416

H −10.6229897 −0.5982142 9.3462273

C −4.0288622 −0.7771437 4.4414573

H −3.0519971 −1.0623361 3.9936804

H −4.3580906 −1.6308385 5.0607686

S −5.3231113 −0.3379424 3.2286377

C −4.3064880 −7.5476450 2.0184130

H −4.5241608 −8.1991813 1.1502046

H −5.2259598 −6.9909574 2.2727992

O −3.2084855 −6.6830079 1.7582894

H −3.5121005 −6.1155568 1.0031016

C −0.5843298 −4.1499681 9.9815016

H −0.1852582 −3.8617743 10.9674760

H −0.3389727 −3.3309095 9.2890948

C −2.1019068 −4.3770630 10.0570458

H −2.4968286 −4.6410050 9.0632345

H −2.3212013 −5.2316497 10.7180726

C −2.9206585 −3.1785420 10.5307630

H −2.5381037 −2.7410739 11.4633731

H −2.9591831 −2.3750113 9.7819662

N −4.3365786 −3.6038641 10.7955134

H −4.9898052 −2.7972865 10.7588930

H −4.6705249 −4.3267210 10.1295153

H −4.4076532 −4.0010363 11.7517938

N −8.2721119 −6.3282526 7.5038966

C −7.3985170 −5.3035209 7.2138047

N −7.0896463 −4.2188151 7.9315008

C −6.1318138 −3.4574102 7.3616528

N −5.6678744 −2.4019678 8.0822049

N −5.5640019 −3.6746995 6.1522760

C −5.9276307 −4.7268920 5.4337406

O −5.4057929 −4.9692818 4.2377200

C −6.8573903 −5.6518136 5.9656051

N −7.3616226 −6.8570387 5.5196479

C −8.1953061 −7.2369725 6.4446178

C −4.6905884 −3.9255069 3.5401648

H −6.1871706 −2.0904972 8.8978506

H −5.0690139 −1.7039815 7.6405460

H −8.8000504 −8.1454823 6.4290783

H −4.3909303 −4.3838018 2.5910402

H −5.3205745 −3.0316029 3.3684557

H −3.7996618 −3.6343780 4.1101216

H −11.3305884 1.1190442 11.7696992

H −3.8232139 0.0589131 5.1098771

H −4.1218493 −8.1945041 2.8760545

H −0.0606374 −5.0414603 9.6364580

H −8.8085016 −6.4414446 8.3402255

#### TS1

C −11.6740957 0.0863987 11.8069719

H −12.0684263 −0.0977585 12.8176234

H −12.5188517 −0.0186267 11.1031566

C −10.5898190 −0.9250489 11.4792514

C −9.9818594 −1.6890818 12.4891452

H −10.3349193 −1.5829549 13.5178760

C −8.9380526 −2.5778761 12.2161293

H −8.4799937 −3.1598537 13.0191202

C −8.4713911 −2.7356033 10.9055981

O −7.4100007 −3.5639609 10.6600961

H −7.4070390 −3.8511529 9.7070092

C −9.1181648 −2.0359304 9.8725922

H −8.8078050 −2.2070596 8.8397224

C −10.1590049 −1.1510199 10.1616563

H −10.6378705 −0.6192411 9.3341889

C −3.5290010 −1.0160003 4.3400012

H −2.4799059 −1.1411771 4.0188371

H −3.7976115 −1.8824931 4.9706334

S −4.6930013 −0.9520003 2.9470008

C −4.2930107 −7.4838219 2.0558821

H −4.5589253 −8.1141643 1.1840559

H −5.1796117 −6.8960423 2.3528485

O −3.1730265 −6.6559353 1.7729713

H −3.4879342 −6.0610212 1.0489830

C −0.5918405 −4.1720212 10.0050960

H −0.2021645 −3.9050344 11.0009403

H −0.3397347 −3.3387677 9.3327203

C −2.1101598 −4.4009320 10.0581687

H −2.4942141 −4.6481757 9.0555544

H −2.3380411 −5.2657729 10.7029838

C −2.9310828 −3.2080469 10.5429773

H −2.5529553 −2.7830135 11.4832121

H −2.9630775 −2.3958194 9.8036861

N −4.3493210 −3.6299928 10.7918960

H −4.9989619 −2.8222198 10.7398826

H −4.6761787 −4.3516047 10.1203039

H −4.4329239 −4.0222616 11.7484687

N −8.2921399 −6.3681734 7.5394404

C −7.3918296 −5.3623867 7.2577328

N −7.0811730 −4.2764329 7.9829808

C −6.1008789 −3.5410677 7.4147129

N −5.6237315 −2.4822908 8.1482096

N −5.5166851 −3.7686409 6.2290048

C −5.8670016 −4.8350014 5.4660015

O −5.3420015 −5.0280014 4.3090012

C −6.8400677 −5.7236720 6.0211169

N −7.3667638 −6.9192685 5.5711994

C −8.2201364 −7.2824517 6.4849046

C −5.0040014 −3.3890010 3.4260010

H −6.1911121 −2.1278903 8.9129131

H −5.0398395 −1.7881254 7.6843063

H −8.8433513 −8.1778511 6.4646073

H −5.4900015 −3.6780010 2.4990007

H −5.5530016 −2.8470008 4.1740012

H −3.9250011 −3.4710010 3.4920010

H −11.3306921 1.1200437 11.7655078

H −3.5671055 −0.1338538 4.9790887

H −4.1113962 −8.1522699 2.8974643

H −0.0627922 −5.0546553 9.6457466

H −8.8307144 −6.4732834 8.3754199

#### IM

C −11.6766312 0.0970488 11.8165739

H −12.0756132 −0.0820295 12.8263494

H −12.5186894 −0.0090981 11.1096451

C −10.5923438 −0.9161350 11.4995671

C −9.9824069 −1.6678473 12.5173810

H −10.3353149 −1.5498424 13.5448433

C −8.9398001 −2.5595570 12.2529402

H −8.4806476 −3.1332429 13.0612539

C −8.4741257 −2.7343254 10.9429679

O −7.4229382 −3.5694936 10.7064620

H −7.3857814 −3.8216793 9.7369181

C −9.1219930 −2.0420480 9.9032211

H −8.8172221 −2.2311180 8.8721318

C −10.1625826 −1.1546730 10.1844491

H −10.6447664 −0.6322065 9.3523107

C −3.7595443 −1.2401504 4.4697248

H −2.7416126 −1.4067353 4.0801501

H −4.0610636 −2.0766446 5.1298065

S −4.9510711 −1.2152150 3.0933754

C −4.2946090 −7.4178051 2.1021909

H −4.6394075 −8.0237674 1.2406214

H −5.1289310 −6.7881028 2.4582892

O −3.1541197 −6.6428004 1.7546992

H −3.4851145 −6.0279208 1.0581305

C −0.5938676 −4.1697154 9.9945715

H −0.1972050 −3.8707857 10.9784797

H −0.3563893 −3.3535425 9.2956437

C −2.1091491 −4.4158889 10.0688150

H −2.4984340 −4.7103723 9.0813666

H −2.3196740 −5.2574859 10.7489873

C −2.9415702 −3.2148409 10.5125568

H −2.5633730 −2.7481327 11.4329889

H −2.9884236 −2.4350816 9.7397695

N −4.3544486 −3.6368745 10.7812856

H −5.0067953 −2.8351644 10.6891078

H −4.6767921 −4.3828465 10.1335820

H −4.4363856 −3.9926906 11.7517259

N −8.2196143 −6.2459535 7.5011199

C −7.3160415 −5.2240502 7.2768335

N −7.0403578 −4.1556447 8.0484571

C −6.0371459 −3.4038436 7.5356437

N −5.6370573 −2.3422476 8.3202933

N −5.3604399 −3.6052233 6.4045351

C −5.6192811 −4.7014731 5.6050270

O −4.9263687 −4.9458544 4.5948078

C −6.6961152 −5.5515241 6.0703917

N −7.1858734 −6.7476223 5.5754017

C −8.0832280 −7.1392680 6.4356420

C −4.4703717 −2.8542445 2.4404953

H −6.2835850 −1.9895468 9.0190677

H −5.0374312 −1.6351358 7.8995730

H −8.6972228 −8.0387808 6.3661723

H −5.1659629 −3.1436112 1.6435052

H −4.5358610 −3.6090457 3.2437878

H −3.4449174 −2.7972696 2.0462882

H −11.3290814 1.1291474 11.7713100

H −3.7474123 −0.3112726 5.0399055

H −4.1026757 −8.1071518 2.9244012

H −0.0587554 −5.0552413 9.6516925

H −8.7705328 −6.3886102 8.3233791

#### TS2-K

C −11.8172852 0.1505567 11.9335395

H −12.2033768 0.0445050 12.9567960

H −12.6796703 0.0634185 11.2486849

C −10.8224170 −0.9419287 11.6520507

C −10.2328090 −1.6749236 12.6936307

H −10.5255169 −1.4566702 13.7211336

C −9.2887411 −2.6689298 12.4480613

H −8.8363127 −3.2326147 13.2657829

C −8.9137892 −2.9640063 11.1315530

O −7.9674483 −3.9097506 10.9158849

H −7.9002961 −4.0681373 9.9478709

C −9.5359201 −2.2696283 10.0737926

H −9.3191952 −2.5481171 9.0426085

C −10.4755771 −1.2773693 10.3405896

H −10.9553287 −0.7638477 9.5018047

C −3.6171482 −1.4228008 4.4720316

H −2.6008074 −1.5066895 4.0526394

H −3.8062081 −2.2840484 5.1361355

S −4.8697615 −1.5048340 3.1524741

C −4.3102294 −7.2730972 2.1840544

H −4.6579491 −7.8711904 1.3197485

H −5.1562509 −6.6568844 2.5394997

O −3.1730841 −6.4923813 1.8416811

H −3.4752486 −5.9545942 1.0693044

C −0.5511894 −4.1881875 10.0485538

H −0.2508995 −3.9910876 11.0902476

H −0.2257825 −3.3186904 9.4550991

C −2.0652219 −4.3957424 9.9217520

H −2.3106593 −4.6715383 8.8862326

H −2.3963725 −5.2430992 10.5453527

C −2.9375683 −3.1872946 10.2392581

H −3.0921828 −3.0856679 11.3267340

H −2.4830993 −2.2428042 9.9009544

N −4.2544968 −3.3317765 9.5729060

H −4.8790393 −2.6350112 10.0001341

H −4.5944905 −3.2912034 8.2038187

H −4.6486720 −4.2589696 9.7986218

N −8.2943727 −6.2305178 7.4115544

C −7.5164821 −5.0973031 7.3709805

N −7.6236940 −3.9993569 8.1401688

C −6.6139909 −3.1477863 7.9746543

N −6.6707068 −1.9483261 8.5846948

N −5.4703715 −3.3947012 7.2641990

C −5.4682560 −4.4003723 6.2791707

O −4.5356938 −4.4948961 5.4679190

C −6.5936063 −5.3076963 6.3542902

N −6.8034675 −6.5453873 5.7649498

C −7.8022264 −7.0782291 6.4174290

C −4.5187884 −3.2244446 2.6392678

H −7.3475144 −1.7925982 9.3263532

H −5.8594434 −1.3370149 8.5520547

H −8.2508200 −8.0550889 6.2308941

H −5.2347395 −3.5016647 1.8556599

H −4.6455637 −3.8899903 3.5044294

H −3.4888810 −3.3121363 2.2632565

H −11.3943566 1.1482844 11.8163304

H −3.6794541 −0.4897529 5.0320376

H −4.1090792 −7.9709371 2.9968430

H −0.0168672 −5.0617177 9.6750411

H −8.8976077 −6.4512616 8.1779433

#### PC-K

C −11.7483551 0.1346660 11.8681535

H −12.1481691 −0.0045432 12.8832378

H −12.5978022 0.0386891 11.1679015

C −10.7098596 −0.9235049 11.5802901

C −10.1508020 −1.6852247 12.6185906

H −10.5003026 −1.5231961 13.6400625

C −9.1640056 −2.6397745 12.3783329

H −8.7346807 −3.2236164 13.1944951

C −8.7075295 −2.8642578 11.0737881

O −7.7200075 −3.7746690 10.8755410

H −7.6412257 −3.9829901 9.9199206

C −9.2927491 −2.1438539 10.0161173

H −8.9850698 −2.3481071 8.9894881

C −10.2805238 −1.1947826 10.2743601

H −10.7224400 −0.6517079 9.4327708

C −3.5846507 −1.3502365 4.2623482

H −2.5940862 −1.3798812 3.7801950

H −3.6932878 −2.2647496 4.8655827

S −4.9306849 −1.3864993 3.0374804

C −4.3053356 −7.3796147 2.1739448

H −4.6357440 −7.9900308 1.3111006

H −5.1650549 −6.7826467 2.5273405

O −3.1884957 −6.5697944 1.8306796

H −3.5167512 −6.0022853 1.0923326

C −0.4645633 −4.1892260 10.1924389

H −0.1464138 −4.1966877 11.2487802

H −0.0724863 −3.2543127 9.7571613

C −1.9905897 −4.2294433 10.0845841

H −2.3032664 −4.1561053 9.0261202

H −2.3868258 −5.1931527 10.4508353

C −2.7094770 −3.1068687 10.8277672

H −2.5721563 −3.2441894 11.9196925

H −2.2445465 −2.1357553 10.5763018

N −4.1062563 −3.0778892 10.4121266

H −4.6091876 −2.3263170 10.8946738

H −4.7723649 −2.9487973 6.5037753

H −4.5663625 −3.9388394 10.7184340

N −8.3050634 −6.2898126 7.4650913

C −7.4622074 −5.2198526 7.2921694

N −7.3338846 −4.1452645 8.0910310

C −6.3527938 −3.3233605 7.7533160

N −6.1371330 −2.1703141 8.3991945

N −5.5502971 −3.5802881 6.6627960

C −5.6436357 −4.6743372 5.7823326

O −4.7952987 −4.8119318 4.8942809

C −6.7402691 −5.5234250 6.1305233

N −7.1249215 −6.7454623 5.6142349

C −8.0496879 −7.1814889 6.4228527

C −4.5323502 −3.0369583 2.3584957

H −6.5048646 −2.0750734 9.3483168

H −5.2076366 −1.7501246 8.3285911

H −8.6032448 −8.1170798 6.3283303

H −5.2357103 −3.2588606 1.5470762

H −4.6427535 −3.7970754 3.1437868

H −3.5022329 −3.0359491 1.9777979

H −11.3610667 1.1503122 11.7872987

H −3.6328047 −0.4620493 4.8923257

H −4.0903715 −8.0688707 2.9905131

H 0.0020915 −5.0334351 9.6848844

H −8.8579320 −6.4577609 8.2812421

#### TS2-Y

C −11.5048719 0.0462201 11.8109031

H −11.8800567 −0.1783765 12.8210958

H −12.3454396 −0.1098285 11.1109014

C −10.3468048 −0.8889280 11.4560862

C −9.7395851 −1.7277379 12.4079848

H −10.1398496 −1.7443980 13.4253338

C −8.6346159 −2.5404874 12.1042044

H −8.2015252 −3.1794144 12.8784430

C −8.0833353 −2.5542786 10.8108724

O −7.0376495 −3.3486070 10.4371680

H −7.1617538 −3.9267285 8.9674402

C −8.7380132 −1.7722255 9.8351337

H −8.3650721 −1.8046503 8.8122608

C −9.8383257 −0.9726192 10.1477915

H −10.2992907 −0.3837002 9.3472693

C −3.6731802 −1.1792897 4.4265788

H −2.6548794 −1.3350807 4.0350875

H −3.9732320 −2.0224057 5.0757564

S −4.8717479 −1.1577609 3.0568755

C −4.2430276 −7.4377012 2.1085550

H −4.5683035 −8.0520882 1.2454278

H −5.0940461 −6.8249864 2.4535923

O −3.1161620 −6.6399685 1.7706557

H −3.4546676 −6.0232938 1.0801181

C −0.8190434 −4.2354654 9.6714116

H −0.2606876 −3.4038578 10.1345725

H −0.9883701 −3.9661635 8.6124720

C −2.1687734 −4.4475815 10.4189682

H −2.5559031 −5.4653593 10.2522447

H −1.9804187 −4.3917342 11.5002119

C −3.3303652 −3.4677465 10.1104014

H −3.0982302 −2.4415413 10.4374843

H −3.5235449 −3.4292466 9.0263897

N −4.5967912 −3.8922511 10.7802989

H −5.8341229 −3.3794266 10.6703707

H −4.7844536 −4.8653503 10.4911057

H −4.4336904 −3.9277471 11.8003767

N −8.1282185 −6.2606085 7.5335462

C −7.2771488 −5.2284800 7.2329085

N −6.9719447 −4.1018482 7.9276486

C −6.0285594 −3.2778357 7.3503142

N −5.7672025 −2.1428638 8.0308826

N −5.4174399 −3.5291344 6.2049607

C −5.6562972 −4.6860875 5.4833813

O −4.9761469 −4.9767686 4.4885793

C −6.7053431 −5.5464638 6.0118275

N −7.1720242 −6.7699687 5.5707426

C −8.0140749 −7.1741346 6.4772121

C −4.3880608 −2.7964041 2.4071189

H −6.0532007 −2.0432197 9.0055961

H −5.0098867 −1.5351509 7.7272388

H −8.6057226 −8.0903895 6.4536676

H −5.0607637 −3.0734564 1.5865949

H −4.4804082 −3.5602392 3.1965279

H −3.3513014 −2.7451018 2.0432626

H −11.2421619 1.1037571 11.7850924

H −3.6674185 −0.2562936 5.0063307

H −4.0456102 −8.1228274 2.9329915

H −0.1708499 −5.1113891 9.6455284

H −8.6694448 −6.3900159 8.3643893

#### PC-Y

C −11.6510545 0.0867775 11.8349306

H −12.0431343 −0.0950714 12.8462784

H −12.4924501 −0.0382191 11.1300677

C −10.5440364 −0.9022793 11.5131909

C −10.0106780 −1.7519697 12.4958741

H −10.4357382 −1.7338659 13.5025599

C −8.9414277 −2.6153646 12.2213410

H −8.5457870 −3.2718617 12.9994991

C −8.3774817 −2.6368447 10.9441605

O −7.3475707 −3.5063217 10.6117885

H −7.2471440 −3.9903454 8.9682193

C −8.9400486 −1.8451487 9.9357371

H −8.5186768 −1.8938737 8.9322098

C −10.0125543 −1.0012658 10.2163439

H −10.4268448 −0.3890546 9.4094735

C −3.7007529 −1.1947160 4.4295759

H −2.6812710 −1.3561117 4.0436902

H −4.0069188 −2.0351112 5.0789913

S −4.8902144 −1.1639676 3.0523208

C −4.2457973 −7.4198416 2.1204228

H −4.5750559 −8.0354807 1.2597357

H −5.0954482 −6.8061523 2.4672655

O −3.1193256 −6.6242838 1.7765048

H −3.4575695 −6.0139548 1.0802131

C −0.7275060 −4.2247493 9.6530526

H −0.1486620 −3.4040469 10.1111515

H −0.9005917 −3.9536716 8.5945269

C −2.0622805 −4.4201949 10.4077133

H −2.4642731 −5.4370341 10.2655029

H −1.8735838 −4.3406112 11.4877384

C −3.2057445 −3.4392999 10.0759032

H −2.9308306 −2.4111880 10.3719855

H −3.3667140 −3.4191247 8.9814825

N −4.4738323 −3.8002902 10.7238365

H −6.4249595 −3.2278569 10.8606250

H −4.7084795 −4.7590412 10.4356565

H −4.3228496 −3.8420286 11.7419414

N −8.1592293 −6.2828727 7.5398173

C −7.3115721 −5.2458461 7.2517732

N −7.0135421 −4.1209349 7.9581379

C −6.0578452 −3.2910861 7.4039464

N −5.8048368 −2.1536936 8.0812340

N −5.4460231 −3.5348009 6.2586937

C −5.6805913 −4.6819268 5.5202577

O −4.9984697 −4.9591138 4.5240612

C −6.7304334 −5.5506958 6.0337308

N −7.1906518 −6.7712208 5.5788739

C −8.0367831 −7.1867631 6.4762184

C −4.4186024 −2.8083924 2.4076517

H −6.0149053 −2.0547697 9.0763975

H −5.0297070 −1.5715724 7.7688413

H −8.6249437 −8.1049230 6.4422457

H −5.0946659 −3.0844410 1.5895525

H −4.5156341 −3.5643636 3.2034269

H −3.3815446 −2.7680791 2.0440181

H −11.3181053 1.1232970 11.7816445

H −3.6930308 −0.2714552 5.0088832

H −4.0457124 −8.1042253 2.9448330

H −0.1081744 −5.1213098 9.6270762

H −8.7112094 −6.4145201 8.3632001

## REFERENCES

(1) Yi, C., & He, C. (2013). DNA Repair by Reversal of DNA Damage. Cold Spring Harbor Perspectives in Biology, 5(1), a012575–a012575.

(2) Daniels, D. S., & Tainer, J. A. (2000). Conserved structural motifs governing the stoichiometric repair of alkylated DNA by O6-alkylguanine-DNA alkyltransferase. DNA Repair, 460(3-4), 151–163.

(3) Daniels, D. S., Woo, T. T., Luu, K. X., Noll, D. M., Clarke, N. D., Pegg, A. E., & Tainer, J. A. (2004). DNA binding and nucleotide flipping by the human DNA repair protein AGT. Nature Structural & Molecular Biology, 11(8), 714–720.

(4) Tardiff, R. G.; Lohman, P. H. M.; Wogan, G. N.; Chemicals, S. G. on M. for the S. E. of. Methods to Assess DNA Damage and Repair: Interspecies Comparisons; Wiley, 1994.

(5) Chatterjee, N., & Walker, G. C. (2017). Mechanisms of DNA damage, repair, and mutagenesis. Environmental and Molecular Mutagenesis, 58(5), 235–263.

(6) Margison, G. P., & Santibáñez-Koref, M. F. (2002). O6-alkylguanine-DNA alkyltransferase: Role in carcinogenesis and chemotherapy. BioEssays, 24(3), 255–266.

(7) Pegg A. E. (1990). Mammalian O6-alkylguanine-DNA alkyltransferase: regulation and importance in response to alkylating carcinogenic and therapeutic agents. Cancer research, 50(19), 6119–6129

(8) Dolan, M. E., Moschel, R. C., & Pegg, A. E. (1990). Depletion of mammalian O6-alkylguanine-DNA alkyltransferase activity by O6-benzylguanine provides a means to evaluate the role of this protein in protection against carcinogenic and therapeutic alkylating agents. Proceedings of the National Academy of Sciences, 87(14), 5368–5372.

(9) Fang, Q., Noronha, A. M., Murphy, S. P., Wilds, C. J., Tubbs, J. L., Tainer, J. A., … Pegg, A. E. (2008). Repair ofO6-G-Alkyl-O6-G Interstrand Cross-Links by HumanO6-Alkylguanine-DNA Alkyltransferase†. Biochemistry, 47(41), 10892–10903.

(10) Wibley, J. E. A. (2000). Crystal structure of the human O6-alkylguanine-DNA alkyltransferase. Nucleic Acids Research, 28(2), 393–401.

(11) Mishina, Y., Duguid, E. M., & He, C. (2006). Direct reversal of DNA alkylation damage. Chemical reviews, 106(2), 215–232.

(12) Duguid, E. M., Rice, P. A., & He, C. (2005). The Structure of the Human AGT Protein Bound to DNA and its Implications for Damage Detection. Journal of Molecular Biology, 350(4), 657–666.

(13) Tubbs, J. L., Pegg, A. E., & Tainer, J. A. (2007). DNA binding, nucleotide flipping, and the helix-turn-helix motif in base repair by O6-alkylguanine-DNA alkyltransferase and its implications for cancer chemotherapy. DNA Repair, 6(8), 1100–1115.

(14) Gerson, S. L., Allay, E., Vitantonio, K., & Dumenco, L. L. (1995). Determinants of O6-alkylguanine-DNA alkyltransferase activity in human colon cancer. Clinical cancer research : an official journal of the American Association for Cancer Research, 1(5), 519–525.

(15) Daniels, D. S., Mol, C. D., Arvai, A. S., Kanugula, S., Pegg, A. E., & Tainer, J. A. (2000). Active and alkylated human AGT structures: a novel zinc site, inhibitor and extrahelical base binding. The EMBO journal, 19(7), 1719–1730.

(16) Rasimas, J. J., Kanugula, S., Dalessio, P. M., Ropson, I. J., Fried, M. G., & Pegg, A. E. (2003). Effects of Zinc Occupancy on HumanO6-Alkylguanine-DNA Alkyltransferase†. Biochemistry, 42(4), 980–990.

(17) Jena, N. R., Shukla, P. K., Jena, H. S., Mishra, P. C., & Suhai, S. (2009). O6-methylguanine repair by O6-alkylguanine-DNA alkyltransferase. The journal of physical chemistry. B, 113(51), 16285–16290.

(18) Hou, Q., Du, L., Gao, J., Liu, Y., & Liu, C. (2010). QM/MM Study on the Reaction Mechanism of O6-Alkylguanine-DNA Alkyltransferase. The Journal of Physical Chemistry B, 114(46), 15296–15300.

(19) Tian, C., Kasavajhala, K., Belfon, K. A. A., Raguette, L., Huang, H., Migues, A. N., Bickel, J., Wang, Y., Pincay, J., Wu, Q., & Simmerling, C. (2019). ff19SB: Amino-Acid-Specific Protein Backbone Parameters Trained against Quantum Mechanics Energy Surfaces in Solution. Journal of Chemical Theory and Computation, 16(1), 528–552.

(20) Izaguirre, J. A., Catarello, D. P., Wozniak, J. M., & Skeel, R. D. (2001). Langevin stabilization of molecular dynamics. The Journal of Chemical Physics, 114(5), 2090–2098.

(21) Berendsen, H. J. C., Postma, J. P. M., van Gunsteren, W. F., DiNola, A., & Haak, J. R. (1984). Molecular dynamics with coupling to an external bath. The Journal of Chemical Physics, 81(8), 3684–3690.

(22) Åqvist, J.; Wennerström, P.; Nervall, M.; Bjelic, S.; Brandsdal, B. O.(2004). Molecular Dynamics Simulations of Water and Biomolecules with a Monte Carlo Constant Pressure Algorithm. Chemical Physics Letters, 384 (4), 288–294.

(23) Ryckaert, J.-P., Ciccotti, G., & Berendsen, H. J. C. (1977). Numerical integration of the cartesian equations of motion of a system with constraints: molecular dynamics of n-alkanes. Journal of Computational Physics, 23(3), 327–341.

(24) Darden, T., York, D., & Pedersen, L. (1993). Particle mesh Ewald: AnN·log(N) method for Ewald sums in large systems. The Journal of Chemical Physics, 98(12), 10089–10092.

(25) Salomon-Ferrer, R., Götz, A. W., Poole, D., Le Grand, S., & Walker, R. C. (2013). Routine Microsecond Molecular Dynamics Simulations with AMBER on GPUs. 2. Explicit Solvent Particle Mesh Ewald. Journal of Chemical Theory and Computation, 9(9), 3878–3888.

(26) Roe, D. R., & Cheatham, T. E. (2013). PTRAJ and CPPTRAJ: Software for Processing and Analysis of Molecular Dynamics Trajectory Data. Journal of Chemical Theory and Computation, 9(7), 3084–3095.

(27) Humphrey, W., Dalke, A., & Schulten, K. (1996). VMD: Visual molecular dynamics. Journal of Molecular Graphics, 14(1), 33–38.

(28) Sur, S., Tiwari, V., et al. (2017) Naphthalenediimide-linked bisbenzimidazole derivatives as telomeric G-quadruplex-stabilizing ligands with improved anticancer activity. ACS Omega, 2 (3), 966–980.

(29) Chaubey, A.K., Dubey, K.D., Ojha, R.P., Stability and free energy calculation of LNA modified quadruplex: a molecular dynamics study. Journal of computer-aided molecular design 26 (3), 289–299.

(30) Sherwood, P.; de Vries, A. H.; et al.(2003) QUASI: A general purpose implementation of the QM/MM approach and its application to problems in catalysis. Journal of Molecular Structure: THEOCHEM, 632(1-3), 1–28.

(31) Metz, S.; Kästner, J.; Sokol, A. A.; Keal, T. W.; Sherwood, P. ChemShell-a Modular Software Package for QM/MM Simulations. Wiley Interdiscip. Rev. Comput. Mol. Sci. 2014, 4 (2), 101–110.

(32) Balasubramani, S. G.; Chen, G. P.; Coriani, S.; Diedenhofen, M.; Frank, M. S.; Franzke, Y. J.; Furche, F.; Grotjahn, R.; Harding, M. E.; Hättig, C.; Hellweg, A.; Helmich-Paris, B.; Holzer, C.; Huniar, U.; Kaupp, M.; Marefat Khah, A.; Karbalaei Khani, S.; Müller, T.; Mack, F.; Nguyen, B. D.; Parker, S. M.; Perlt, E.; Rappoport, D.; Reiter, K.; Roy, S.; Rückert, M.; Schmitz, G.; Sierka, M.; Tapavicza, E.; Tew, D. P.; Van Wüllen, C.; Voora, V. K.; Weigend, F.; Wodyński, A.; Yu, J. M. (2020). TURBOMOLE: Modular program suite for ab initio quantum-chemical and condensed-matter simulations. The Journal of chemical physics, 152(18), 184107.

(33) Ahlrichs, R., Bär, M., Häser, M., Horn, H., & Kölmel, C. (1989). Electronic structure calculations on workstation computers: The program system turbomole. Chemical Physics Letters, 162(3), 165–169.

(34) Smith, W., & Forester, T. R. (1996). DL_POLY_2.0: A general-purpose parallel molecular dynamics simulation package. Journal of Molecular Graphics, 14(3), 136–141.

(35) Kästner, J., Carr, J. M., Keal, T. W., Thiel, W., Wander, A., & Sherwood, P. (2009). DL-FIND: An Open-Source Geometry Optimizer for Atomistic Simulations. The Journal of Physical Chemistry A, 113(43), 11856–11865.

